# eQTpLot: a user-friendly R package for the visualization and colocalization of eQTL and GWAS signals

**DOI:** 10.1101/2020.08.26.268268

**Authors:** Theodore G. Drivas, Anastasia Lucas, Marylyn D. Ritchie

**Affiliations:** Division of Human Genetics, Children’s Hospital of Philadelphia, Philadelphia, PA, USA; Department of Genetics, Perelman School of Medicine, University of Pennsylvania, Philadelphia, PA, USA; Institute for Biomedical Informatics, Perelman School of Medicine, University of Pennsylvania, Philadelphia, PA, USA

## Abstract

Genomic studies increasingly integrate expression quantitative trait loci (eQTL) information into their analysis pipelines, but few tools exist for the visualization of colocalization between eQTL and GWAS results. Those tools that do exist are limited in their analysis options, and do not integrate eQTL and GWAS information into a single figure panel, making the visualization of colocalization difficult.

To address this issue, we developed the intuitive and user-friendly R package eQTpLot. eQTpLot takes as input standard GWAS and eQTL summary statistics, and optional pairwise LD information, to generate a series of plots visualizing colocalization, correlation, and enrichment between eQTL and GWAS signals for a given gene-trait pair. With eQTpLot, investigators can easily generate a series of customizable plots clearly illustrating, for a given gene-trait pair: 1) colocalization between GWAS and eQTL signals, 2) correlation between GWAS and eQTL p-values, 3) enrichment of eQTLs among trait-significant variants, 4) the LD landscape of the locus in question, and 5) the relationship between the direction of effect of eQTL signals and the direction of effect of colocalizing GWAS peaks. These clear and comprehensive plots provide a unique view of eQTL-GWAS colocalization, allowing for a more complete understanding of the interaction between gene expression and trait associations.

In summary, eQTpLot provides a unique, user-friendly, and intuitive means of visualizing eQTL and GWAS signal colocalization, incorporating novel features not found in other eQTL visualization software. We believe eQTpLot will prove a useful tool for investigators seeking a convenient and customizable visualization of eQTL and GWAS data colocalization.

**Availability and Implementation:** the eQTpLot R package and tutorial are available at https://github.com/RitchieLab/eQTpLot

## Background

Non-protein-coding genetic variants make up the majority of statistically significant associations identified by genome wide association studies (GWAS). As these variants typically do not have obvious consequences for gene function, it can be difficult to map their effects to specific genes. To address this issue, genomic studies have increasingly begun to integrate expression quantitative trait loci (eQTL) information into their analysis pipelines, with the thought that non-coding variants might be exerting their effects on patient phenotypes through the modulation of expression levels of nearby genes. Through this approach, indirect evidence for causality can be obtained if a genetic locus significantly associated with candidate gene expression levels is found to colocalize with a genetic locus significantly associated with the phenotype of interest.

A number of excellent tools have been developed to discover and analyze colocalization between eQTL and GWAS association signals (1–8), but few packages provide the necessary tools to visualize these colocalizations in an intuitive and informative way. LocusCompare (8) allows for the side-by-side visualization of eQTL and GWAS signal colocalization, but does not visually integrate this data. LocusZoom (9) produces a single plot integrating linkage disequilibrium (LD) information and GWAS data, but does not consider eQTL data. Furthermore, no colocalization visualization tool exists that takes into account the direction of effect of an eQTL with relation to the direction of effect of colocalizing GWAS signals.

For these reasons, we developed eQTpLot, an R package for the intuitive visualization of colocalization between eQTL and GWAS signals. In its most basic implementation, eQTpLot takes standard GWAS summary data, formatted as one might obtain from a GWAS analysis in PLINK (10), and eQTL data, formatted as one might download directly from the GTEx portal (11), to generate a series of customizable plots clearly illustrating, for a given gene-trait pair: 1) colocalization between GWAS and eQTL signals, 2) correlation between GWAS and eQTL p-values, 3) enrichment of eQTLs among trait-significant variants, 4) the LD landscape of the locus in question, and 5) the relationship between the directions of effect of eQTL signals and colocalizing GWAS peaks. These clear and comprehensive plots provide a unique view of eQTL-GWAS colocalization, allowing for a more complete understanding of the interaction between gene expression and trait associations. We believe eQTpLot will prove a useful tool for investigators seeking a convenient and robust visualization of genomic data colocalization.

## Implementation

eQTpLot was developed in R and depends on a number of packages for various aspects of its implementation (biomaRt, dplyr, GenomicRanges, ggnewscale, ggplot2, ggplotfy, ggpubr, gridExtra, Gviz, LDheatmap, patchwork) (12–21). The software is freely available on GitHub (https://github.com/RitchieLab/eQTpLot) and can be downloaded for use at the command line, or in any R-based integrated development environment, such as RStudio. Example data and a complete tutorial on the use of eQTpLot and its various features have also been made available on GitHub.

At a minimum, eQTpLot requires two input files, imported into R as data frames: one of GWAS summary statistics (as might be obtained from a standard associations study as completed in PLINK (10)) and one of eQTL summary statistics (as might be downloaded directly from the GTEx portal at gtexportal.org (11)). Table 1 summarizes the formatting parameters of the two required input files and of the two optional input files. Additionally, there are many options that can be specified to generate variations of the main eQTpLot, as discussed below. Table 2 shows the complete list of command line arguments that can be passed to eQTpLot, with descriptions of their use.

**Table 1.** Description of required and optional input data frames for eQTpLot.

|  |  |  |
| --- | --- | --- |
| Required Input Data Frames |  |  |
| **GWAS.df**, a data frame, one row per SNP, with columns as one might obtain from a genome-wide association study performed in PLINK using either the `--logistic` or `--linear` flags |  |  |
| Column Name | Data type | Description |
| CHR | Integer | Chromosome for SNP (sex chromosomes coded numerically) |
| BP | Integer | Chromosomal position for each SNP, in base pairs |
| SNP | Character | Variant ID (such as dbSNP ID "rs...". (*Note: naming scheme must be the same as what is used in the **eQTL.df** to ensure proper SNP matching*) |
| P | Numeric | P-value for the SNP from GWAS analysis |
| BETA | Numeric | Beta for SNP from GWAS analysis |
| PHE<br>(Optional) | Character | Name of the phenotype for which the GWAS data refers. This column is optional and is useful if your **GWAS.df** contains data for multiple phenotypes, such as one might obtain from a PheWAS. If **GWAS.df** does not contain a "PHE" column, eQTpLot will assume all the supplied GWAS data is for a single phenotype, with a name to be specified with the "trait" argument. |
| **eQTL.df**, a data frame, one row per SNP, with columns as one might download directly from the GTEx Portal in .csv format |  |  |
| Column Name | Data type | Description |
| SNP.Id | Character | Variant ID (such as dbSNP ID "rs...". (*Note: naming scheme must be the same as what is used in the **GWAS.df** to ensure proper matching*). |
| Gene.Symbol | Character | Gene symbol to which the eQTL expression data refers (*Note: gene symbol must match entries in **Genes.df** to ensure proper matching*) |
| P.value | Numeric | P-value for the SNP from eQTL analysis |
| NES | Numeric | Normalized effect size for the SNP from eQTL analysis (Per GTEx, defined as the slope of the linear regression, and is computed as the effect of the alternative allele relative to the reference allele in the human genome reference. |
| Tissue | Character | Tissue type to which the eQTL pvalue/NES refer (*Note: **eQTL.df** can contain multiple tissue types*) |

| Optional Input Data Frames |  |  |
| --- | --- | --- |
| **Genes.df**, an optional data frame, one row per gene, with the following columns (*Note: eQTLplot automatically loads a default **Genes.df** containing information for most protein-coding genes for genomic builds hg19 and hg38, but you may wish to specify our own **Genes.df** data frame if your gene of interest is not included in the default data frame, or if your eQTL data uses a different gene naming scheme (for example, Gencode ID instead of gene symbol)*) |  |  |
| Column Name | Data type | Description |
| **Gene** | Character | Gene symbol/name (*Note: gene naming scheme must match entries in **eQTL.df** to ensure proper matching*) |
| **CHR** | Integer | Chromosome the gene is on (*Note: do not include a "chr" prefix, and sex chromosomes should be coded numerically*) |
| **Start** | Integer | Base pair coordinate of the beginning of the gene (*Note: this should be the smaller of the two values between **Start** and **Stop***) |
| **Stop** | Integer | Base pair coordinate of the end of the gene (*Note: this should be the larger of the two values between **Start** and **Stop***) |
| **Build** | Character, "hg19" or "hg38" | The genome build (either hg19 or hg38) for the location data |
| **LD.df**, an optional data frame of SNP linkage data, one row per SNP pair, with columns as one might obtain from a PLINK linkage disequilibrium analysis using the PLINK --r2 option. (*Note: If no **LD.df** is supplied, eQTLplot will plot data without LD information*) |  |  |
| Column Name | Data type | Description |
| **BP_A** | Integer | Base pair position of the first variant in the LD pair |
| **SNP_A** | Character | Variant ID of the first variant in the LD pair (*Note: only variants that also appear in the **GWAS.df** SNP column will be used for LD analysis*) |
| **BP_B** | Integer | Base pair position of the second variant in the LD pair |
| **SNP_B** | Character | Variant ID of the second variant in the LD pair (*Note: only SNPs that also appear in the **GWAS.df** SNP column will be used for LD analysis*) |
| **R2** | Numeric | Squared correlation measure of linkage between the two variants |

**Table 2.**
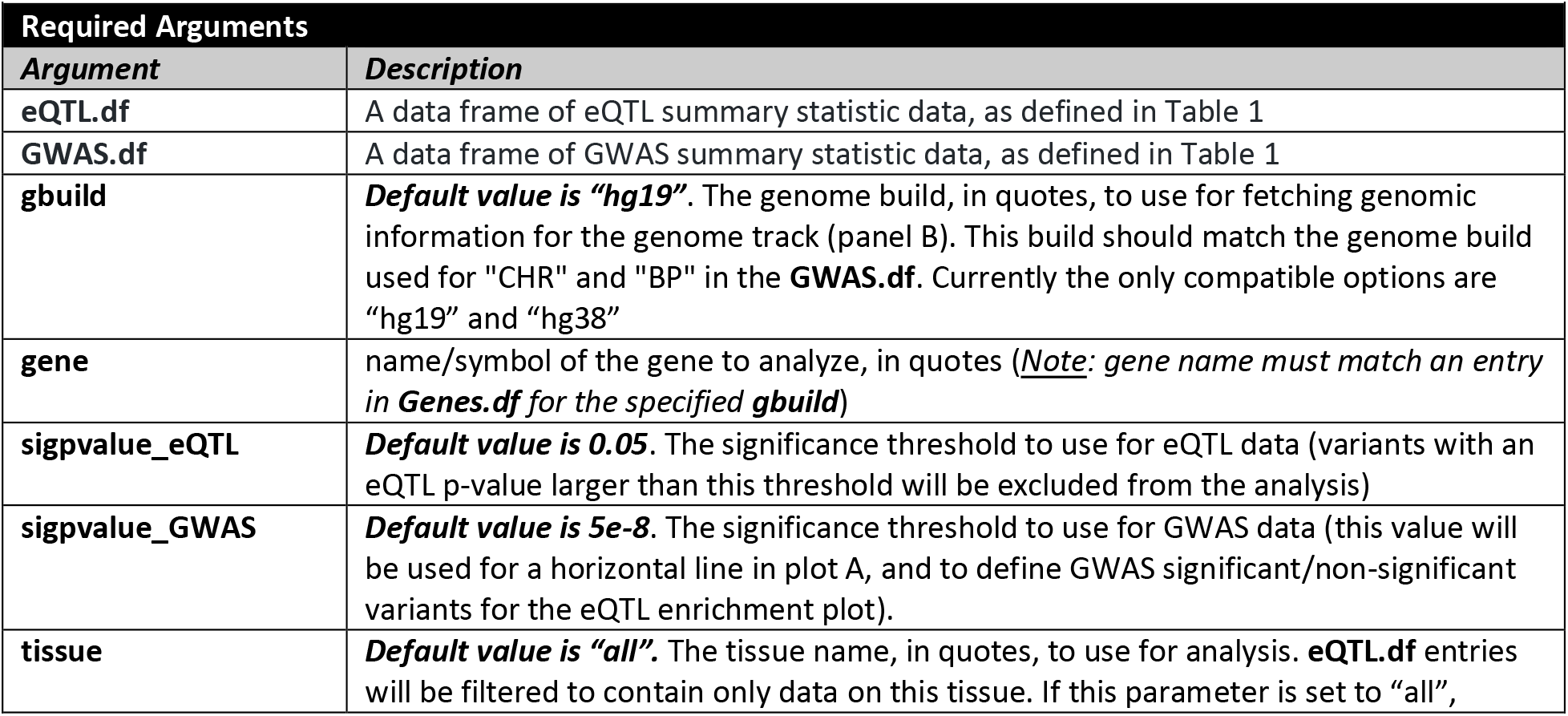
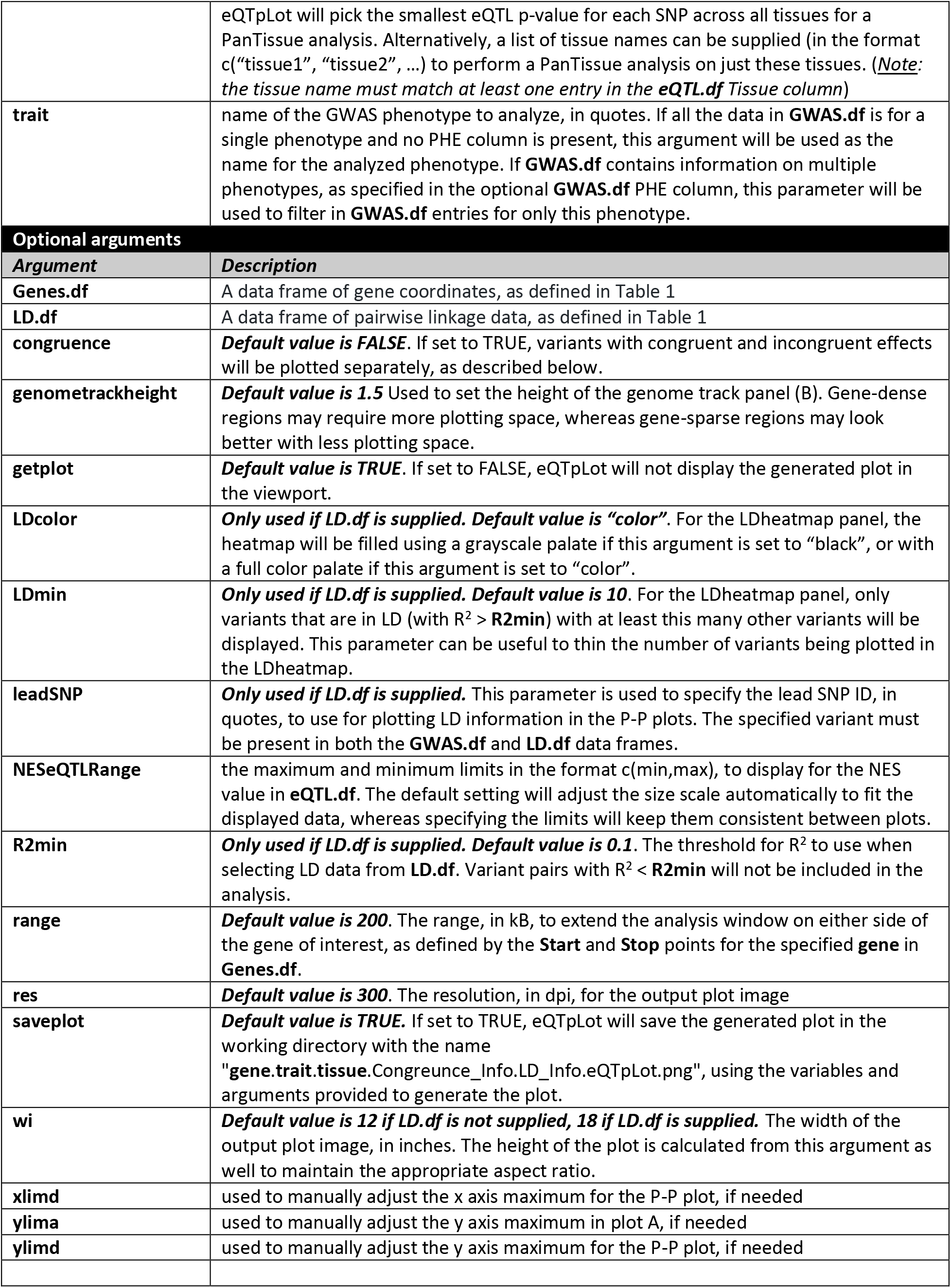
Description of required and optional arguments for eQTpLot.

## Results and Discussion

In its simplest implementation, eQTplot takes as input two data frames, one of GWAS summary data and the other of eQTL summary data, with the user specifying the name of the gene to be analyzed, the GWAS trait to be analyzed (useful if the GWAS data contains information on multiple associations, as one might obtain from a Phenome-wide Association Study (PheWAS)), and the tissue type to use for the eQTL analysis. Using these inputs, eQTpLot generates a series of plots intuitively illustrating the colocalization of GWAS and eQTL signals in chromosomal space, and the enrichment of and correlation between the candidate gene eQTLs and trait-significant variants. Additional parameters and data can be supplied, such as pairwise variant LD information, allowing for an even more comprehensive visualization of the interaction between eQTL and GWAS data within a given genomic locus.

One major implementation feature that sets eQTpLot apart from other eQTL visualization software is the option to divide eQTL/GWAS variants into groups based on their directions of effect. If the option **congruence** is set to TRUE, all variants are divided into two groups: congruous, or those with the same direction of effect on gene expression and the GWAS trait (e.g., a variant that is associated with increased expression of the candidate gene and an increase in the GWAS trait), and incongruous, or those with opposite directions of effect on gene expression and the GWAS trait (e.g., a variant that is associated with increased expression of the candidate gene but a decrease in the GWAS trait). The division between congruous and incongruous variants provides a more nuanced view of the relationship between gene expression level and GWAS associations – a variant associated with increased expression of a candidate gene and an increase in a given GWAS trait would seem to be operating through different mechanisms that a variant that is similarly associated with increased expression of the same candidate gene, but a decrease in the same GWAS trait. eQTpLot intuitively visualizes these differences as described below.

Another important feature of eQTpLot that is not found in other eQTL visualization software is the ability to specify a PanTissue eQTL analysis. In some instances, it may be of interest to visualize a variant’s effect on candidate gene expression across multiple tissue types, or even across all tissues. Such analyses can be accomplished by setting the argument **tissue** to a list of tissues contained within **eQTL.df** (e.g. c(“Adipose_Subcutaneous”, “Adipose_Visceral”)) for a MultiTissue analysis, or by setting the argument **tissue** to “all” for a PanTissue analysis. In a PanTissue analysis, eQTL data across all tissues contained in **eQTL.df** will be collapsed, by variant, into a single pan-tissue eQTL; that is, for each variant, the tissue with the smallest eQTL p-value will be selected. In this way, each variant’s most significant eQTL effect, agnostic of tissue, can be visualized. A similar approach is used in a MultiTissue analysis, but in this case eQTL data will be collapsed, by variant, only across the specified tissues.

What follows is a description of the process used to generate each of the plots produced by eQTpLot, along with a series of use examples to both demonstrate the utility of eQTpLot, and to highlight some of the many options that can be combined to generate different outputs. For these examples we have analyzed a subset of data from our recently-published analysis of quantitative laboratory traits in the UK Biobank (22) – these summary statistics are available in full at https://ritchielab.org/publications/supplementary-data/ajhg-cilium, and the subset of summary data used for our example analyses can be downloaded from the eQTpLot GitHub page such that the reader may experiment with eQTpLot with the pre-supplied data.

### Generation of the main eQTL-GWAS Colocalization Plot

To generate the main eQTL-GWAS Colocalization Plot (Figures 1A, 2A, 3A, 4A), a locus of interest (LOI) is defined to include the target gene’s chromosomal coordinates (as listed in **Genes.df**, for the indicated **gbuild**, for the user-specified **gene**), along with a range of flanking genome (specified with the argument **range**, with a default value of 200 kilobases on either side of the gene). GWAS summary statistics from **GWAS.df** are filtered to include only variants that fall within the LOI. The variants are then plotted in chromosomal space along the horizontal axis, with the inverse log of the p-value of association with the specified GWAS **trait** (p_trait_) plotted along the vertical axis, as one would plot a standard GWAS Manhattan plot. The GWAS significance threshold, **sigpvalue_GWAS** (default value 5e-8), is depicted with a red horizontal line.

**Figure 1.**
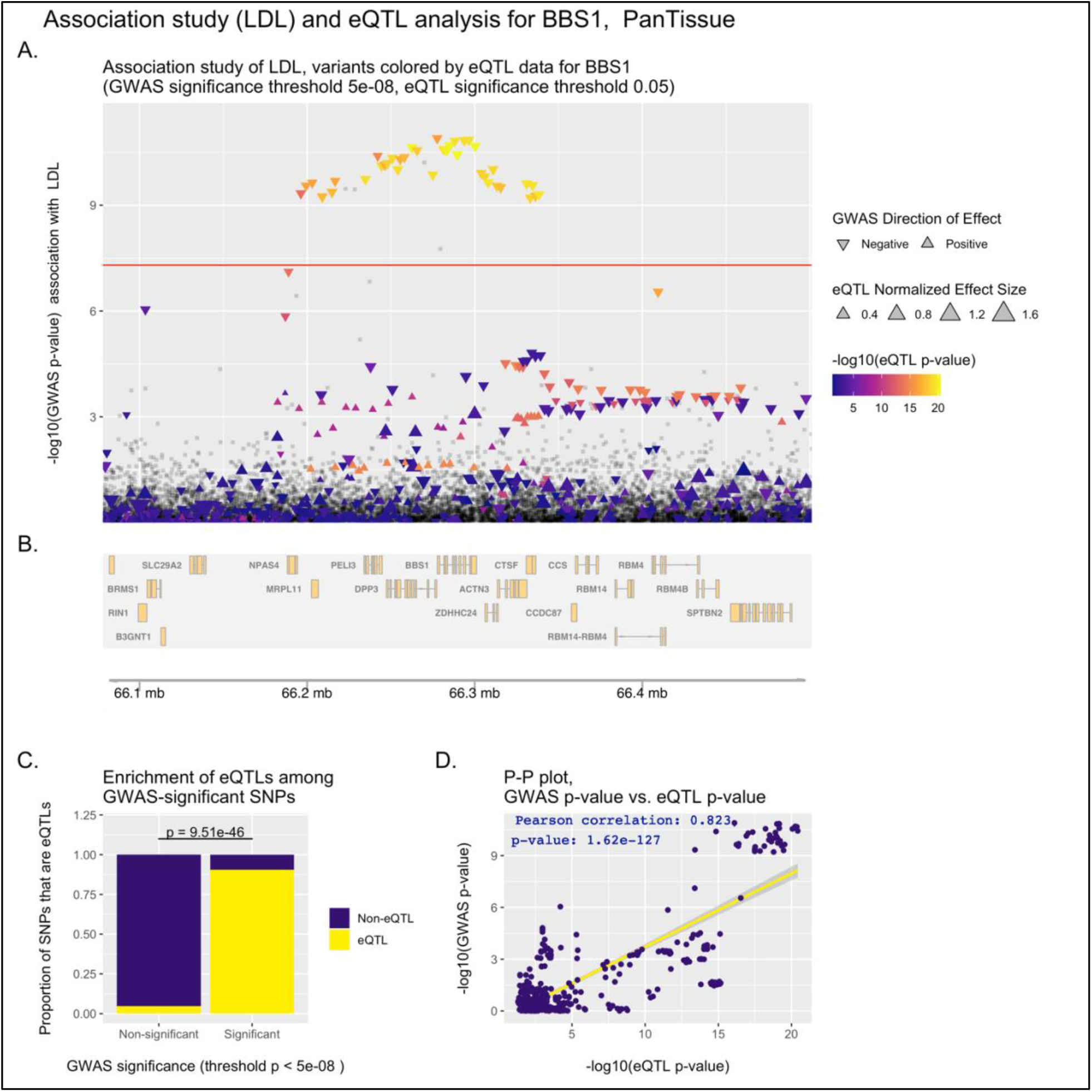
Example eQTpLot illustrating colocalization between variants significantly associated with LDL cholesterol and variants significantly associated with *BBS1* expression levels.

**Figure 2.**
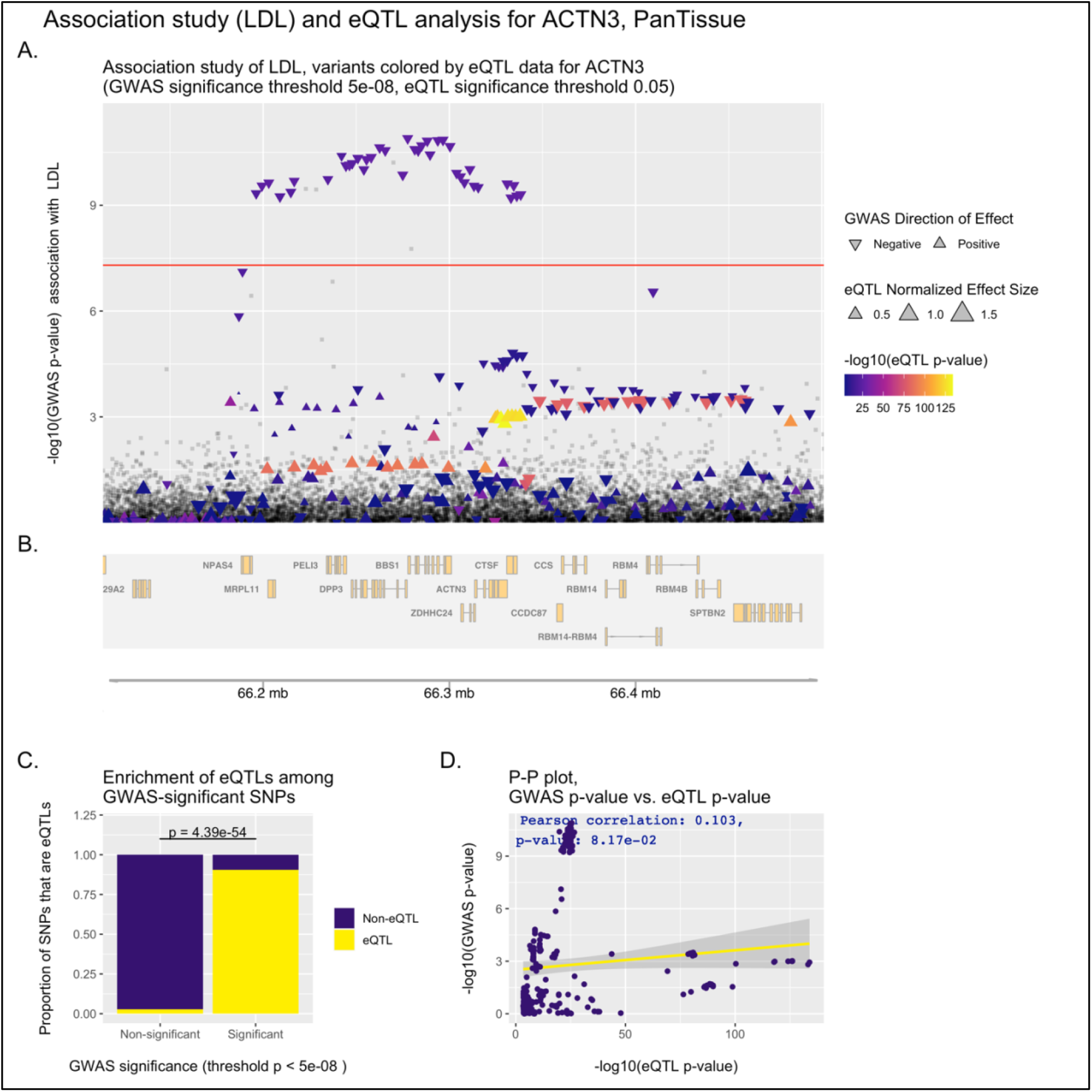
Example eQTpLot illustrating colocalization between variants significantly associated with LDL cholesterol and variants significantly associated with *ACTN3* expression levels.

**Figure 3.**
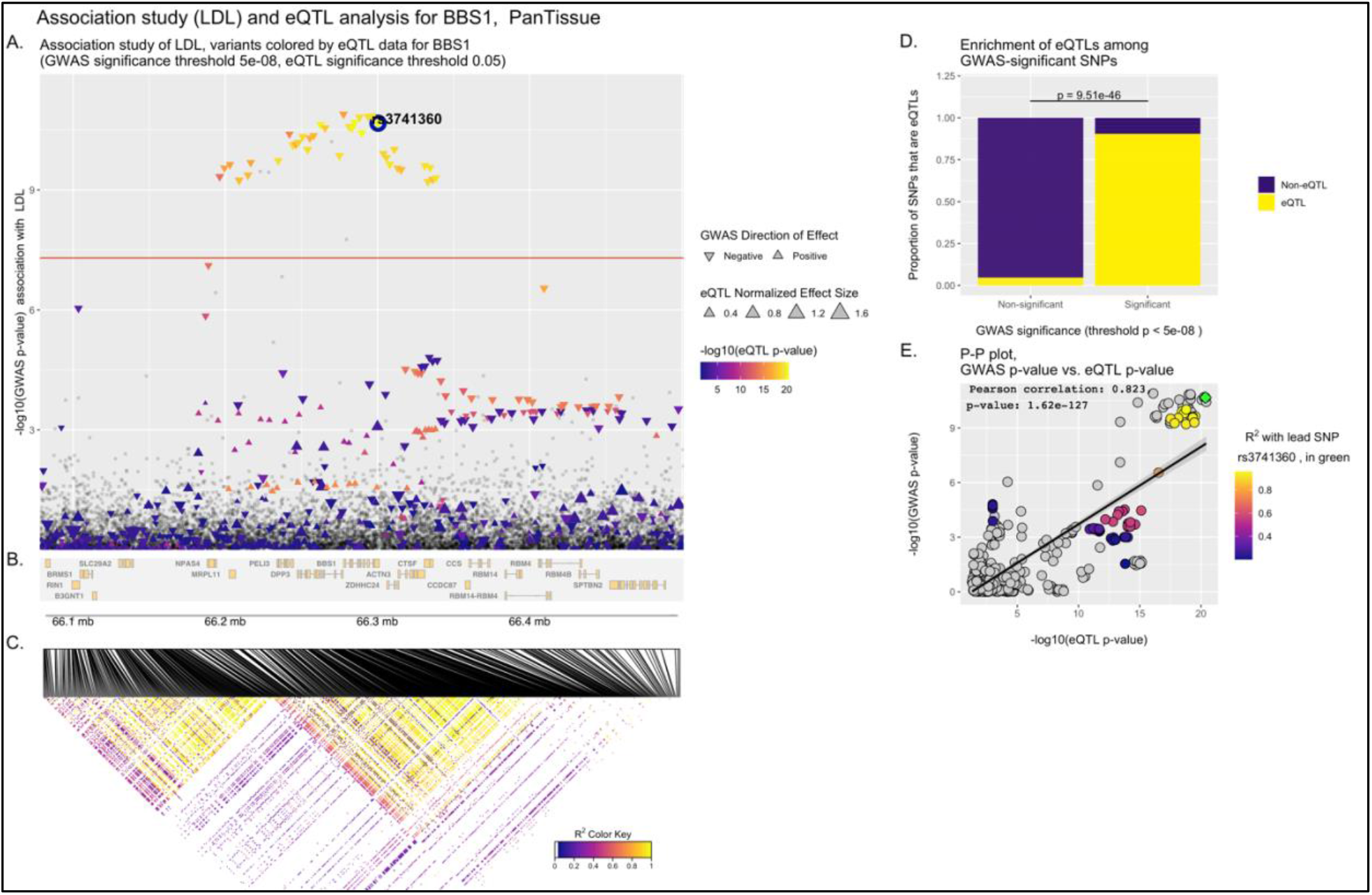
Example eQTpLot illustrating colocalization between variants significantly associated with LDL cholesterol and variants significantly associated with *BBS1* expression levels, with LD information included.

**Figure 4.**
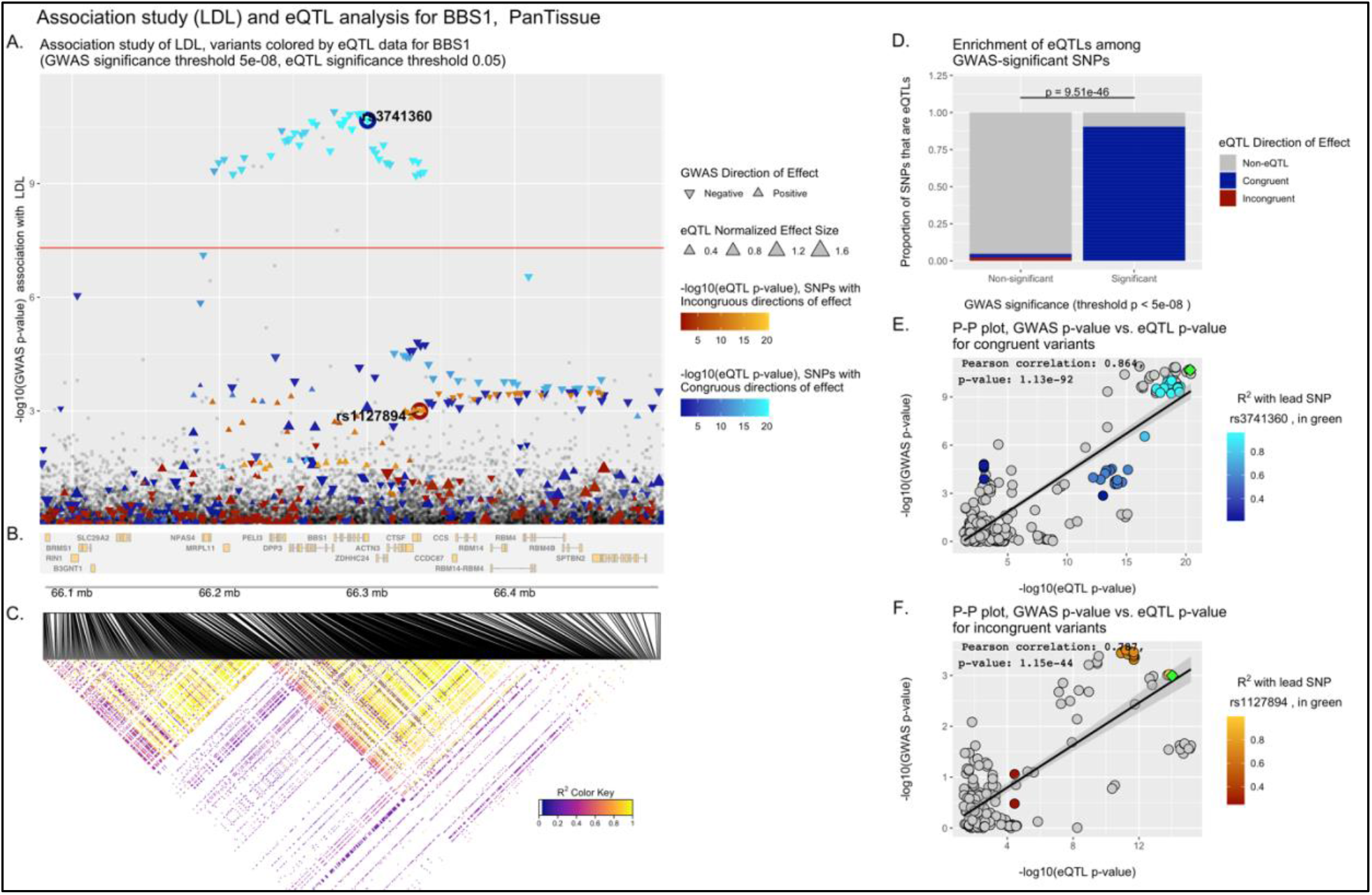
Example eQTpLot illustrating colocalization between variants significantly associated with LDL cholesterol and variants significantly associated with *BBS1* expression levels, with LD information included, and congruent and incongruent variants analyzed separately.

Within this plot, variants that lack eQTL data for the target gene in **eQTL.df** (or for which the eQTL p-value (p_eQTL_) does not meet the specified significance threshold, **sigpvalue_eQTL** (default value 0.05)) are plotted as grey squares. On the other hand, variants that act as eQTLs for the target gene (with p_eQTL_ < **sigpvalue_eQTL**) are plotted as colored triangles, with a color gradient corresponding to the inverse magnitude of p_eQTL_. As noted above, an analysis can be specified to differentiate between variants with congruous versus incongruous effects on the GWAS trait and candidate gene expression levels – if this is the case, variants with congruous effects will be plotted using a blue color scale, while variants with incongruous effects will be plotted using a red color scale (as seen in Figure 4A).The size of each triangle corresponds to the eQTL normalized effect size (NES) for each variant, while the directionality of each triangle is set to correspond to the direction of effect for the variant on the GWAS trait.

A depiction of the genomic positions of all genes within the LOI is generated below the plot using the package Gviz (Figures 1B, 2B, 3B, 4B). (12) If LD data is supplied, in the form of **LD.df**, a third panel illustrating the LD landscape of eQTL variants within the LOI is generated using the package LDheatmap (Figure 3C, 4C). (20) To generate this panel, **LD.df** is filtered to contain only eQTL variants that appear in the plotted LOI, and to include only variant pairs that are in LD with each other with R^2^ > **R2min** (default value of 0.1). This dataset is further filtered to include only variants that are in LD (with R^2^ > **R2min**) with at least a certain number of other variants (user-defined with the argument **LDmin**, default value of 10). These filtering steps are useful in paring down the number of variants to be plotted in the LDheatmap, keeping the most informative variants and reducing the time needed to generate the eQTpLot. A heatmap illustrating the pairwise linkage disequilibrium of the final filtered variant set is subsequently generated below the main eQTL-GWAS Colocalization Plot, with a fill scale corresponding to R^2^ for each variant pair. The location of each variant in chromosomal space is indicated at the top of the heatmap, using the same chromosomal coordinates as displayed in panels A and B.

### Generation of the eQTL Enrichment Plot

For variants within the LOI with p_trait_ less than the specified GWAS significance threshold, **sigpvalue_GWAS**, the proportion that are also eQTLs for the gene of interest (with p_eQTL_ < **sigpvalue_eQTL**) are calculated and plotted, and the same is done for variants with p_trait_ > **sigpvalue_GWAS**, (Figure 1C, 2C, 3D, 4D). Enrichment of candidate gene eQTLs among GWAS-significant variants is determined by Fisher’s exact test. If an analysis differentiating between congruous and incongruous variants is specified, these are considered separately in the analysis (as seen in figure 4D).

### Generation of P-P Correlation Plots

To visualize correlation between p_trait_ and p_eQTL_, each variant within the LOI is plotted with p_eQTL_ along the horizontal axis, and p_trait_ along the vertical axis. Correlation between the two probabilities is visualized by plotting a best-fit linear regression over the points. The Pearson correlation coefficient and p-value of correlation are computed and displayed on the plot as well (Figure 1D, 2D). If an analysis differentiating between congruous and incongruous variants is specified, separate plots are made for each set of variants and superimposed over each other as a single plot, with linear regression lines/Pearson coefficients displayed for both sets.

If LD data is supplied in the form of **LD.df**, a similar plot is generated, but the fill color of each point is set to correspond to the LD R^2^ value for each variant with a specified lead variant, plotted as a green diamond (Figure 3E). This lead variant can be user-specified with the argument **leadSNP** or is otherwise automatically defined as the upper-right-most variant in the P-P plot. This same lead variant is also labelled in the main eQTpLot panel A (Figure 3A). In the case where LD data is provided and an analysis differentiating between congruous and incongruous variants is specified, two separate plots are generated: one for congruous and one for incongruous variants (Figure 4E-F). In each plot, the fill color of each point is set to correspond to the LD R^2^ value for each variant with the lead variant for that specific plot (again defined as the upper-right most variant of the P-P plot), with both the congruous and incongruous lead variants labelled in the main eQTpLot panel A (Figure 4A).

### Use Examples

To more clearly illustrate the use and utility of the eQTpLot software, the following 3 examples are provided. In example 1, the basic implementation of eQTpLot identifies a plausible candidate gene, *BBS1*, for a GWAS association peak for LDL cholesterol on chromosome 11, while also suggesting that a different gene at the same locus, *ACTN3*, is a less plausible based on eQTL colocalization. In example 2 the *BBS1* gene is further investigated through the inclusion of LD data into the eQTpLot analysis. Lastly, in example 3, the analysis is further refined by differentiating between variants with congruous and incongruous directions of effect on *BBS1* expression levels and the LDL cholesterol trait.

#### Example 1 – comparing eQTpLots for two genes within a linkage peak

A GWAS study of LDL cholesterol levels has identified a significant association with a genomic locus at chr11:66,196,265-66,338,300 (build hg19), which contains a number of plausible candidate genes, including *BBS1* and *ACTN3*. eQTpLot is employed, in R, to investigate eQTL colocalization for the *BBS1* gene and the LDL cholesterol signal as follows:

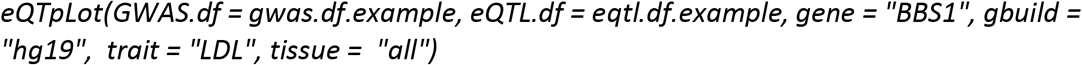

As written, this command will analyze the GWAS data, as contained within GWAS.df.example, within a default 200kb range surrounding the *BBS1* gene, using the preloaded **Genes.df** to define the genomic boundaries of *BBS1* based on genome build hg19. eQTL data from eQTL.df.example will be filtered to contain only data pertaining to *BBS1*. Since **tissue** is set to “all,” eQTpLot will perform a PanTissue analysis, as described above.

The resulting plot (Figure 1) illustrates clear evidence of colocalization between the LDL-significant locus and *BBS1* eQTLs. In Figure 1A, it is easy to see that all variants significantly associated with LDL cholesterol (those plotted above the horizontal red line) are also very significantly associated with *BBS1* expression levels, as indicated by their coloration in bright orange. Figure 1C shows that there is a significant enrichment (p = 9.5e-46 by Fisher’s exact test) for *BBS1* eQTLs among GWAS-significant variants. Lastly, Figure 1D illustrates strong evidence for correlation between p_trait_ and p_eQTL_ for the analyzed variants, with a Pearson correlation coefficient of 0.823 and a p-value of correlation of 1.62e-127 (as displayed on the plot). Taken together, this analysis provides strong evidence for colocalization between variants associated with LDL cholesterol levels and variants associated with *BBS1* expression levels at this genomic locus.

To investigate the possibility that the LDL association signal might also be acting through modulation of the expression of other genes at this locus, the same analysis can be performed, substituting the gene *ACTN3* for the gene *BBS1*, as in the following command:

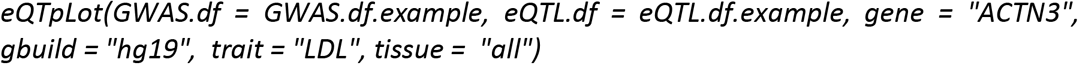

Unlike the previous example for *BBS1*, the resultant plot (Figure 2) shows very poor evidence for colocalization between *ACTN3* eQTLs and LDL cholesterol-significant variants. Although there is significant enrichment for *ACTN3* eQTLs among GWAS-significant variants (Figure 2B), there is poor evidence for correlation between p_trait_ and p_eQTL_ (Figure 2D), and it is intuitively clear in Figure 2A that the eQTL and GWAS signals do not colocalize (the brightest colored points with the strongest association with *ACTN3* expression are not among the variants most significantly associated with LDL cholesterol levels).

#### Example 2 – adding LD information to eQTpLot

To further explore the colocalization between *BBS1* eQTLs and the GWAS association peak for LDL cholesterol, the user can supply LD data to eQTpLot using the argument **LD.df**:

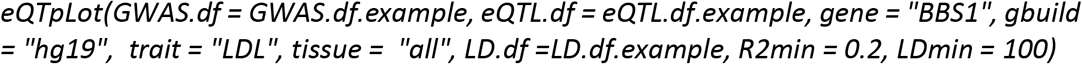

Here the argument **LD.df** refers to the LD.df.example data frame containing a list of pairwise LD correlation measurements between all the variants within the LOI, as one might obtain from a PLINK linkage disequilibrium analysis using the --r2 option. (10) Additionally, the parameter **R2min** is set to 0.2, indicating that **LD.df** should be filtered to drop variant pairs in LD with R^2^ less than 0.2, and **LDmin** is set to 100, indicating that only variants in LD with at least 100 other variants should be plotted in the LD heatmap.

The resultant plot, Figure 3, is different than Figure 1 (the same eQTpLot analysis carried out without LD information) in two important ways. First, a heat map of the LD landscape for all *BBS1* eQTL variants within the LOI is shown in Figure 3C; this heatmap makes it clear that a number of *BBS1* eQTL variants are in strong LD with each other at this locus. Second, the P-P plot, Figure 3E, now includes LD information for all plotted variants; a lead variant, rs3741360, has been defined (by default the upper-right most variant on the P-P plot), and all other variants are plotted with a color scale corresponding to their squared coefficient of linkage correlation with this lead variant. eQTpLot also labels the lead variant in Figure 3A for reference. With the incorporation of this new data, we can now see that most, but not all, of the GWAS-significant variants are in strong LD with each other. This implies that there are at least two distinct LD blocks at the *BBS1* locus with strong evidence of colocalization between the *BBS1* eQTL and LDL GWAS signals.

#### Example 3 – separating congruous from incongruous variants

In addition to including LD data in our eQTpLot analysis, we can also include information on the directions of effect of each variant, with respect to the GWAS trait and *BBS1* expression levels. This is accomplished by setting the argument **congruence** to TRUE:

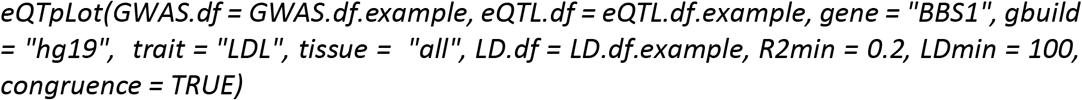

The resulting plot, Figure 4, divides all *BBS1* eQTL variants into two groups: congruent – those variants associated with either an increase in both, or decrease in both *BBS1* expression levels and LDL levels – and incongruent – those variants with opposite directions of effect on *BBS1* expression levels and LDL levels. In carrying out such an analysis, it becomes clear that it is specifically variants with congruent directions of effect that are driving the signal colocalization; that is, variants associated with decreases in *BBS1* expression strongly colocalize with variants associated with decreases in LDL cholesterol.

## Conclusions

eQTpLot provides a unique, user-friendly, and intuitive means of visualizing eQTL and GWAS signal colocalization in a single figure. As plotted by eQTpLot, colocalization between GWAS and eQTL data for a given gene-trait pair is immediately visually obvious, and can be compared across candidate genes to quickly generate hypotheses about the underlying causal mechanisms driving GWAS association peaks. Additionally, eQTpLot allows for Pan- and MultiTissue eQTL analysis, and for the differentiation between eQTL variants with congruous and incongruous directions of effect on GWAS traits – two features not found in any other visualization software. We believe eQTpLot will prove a useful tool for investigators seeking a convenient and customizable visualization of eQTL and GWAS data colocalization.

